# The *Pseudomonas aeruginosa* PSL polysaccharide is a social but non-cheatable trait in biofilms

**DOI:** 10.1101/049783

**Authors:** Yasuhiko Irie, Aled E. L. Roberts, Kasper N. Kragh, Vernita D. Gordon, Jaime Hutchison, Rosalind J. Allen, Gavin Melaugh, Thomas Bjarnsholt, Stuart A. West, Stephen P. Diggle

## Abstract

Extracellular polysaccharides are compounds secreted by microorganisms into the surrounding environment and which are important for surface attachment and maintaining structural integrity within biofilms. The social nature of many extracellular polysaccharides remains unclear, and it has been suggested that they could function as either co-operative public goods, or as traits that provide a competitive advantage. Here we empirically test the co-operative nature of the PSL polysaccharide, which is crucial for the formation of biofilms in *Pseudomonas aeruginosa*. We show that: (1) PSL is not metabolically costly to produce; (2) PSL provides population level benefits in biofilms, for both growth and antibiotic tolerance; (3) the benefits of PSL production are social and are shared with other cells; (4) the benefits of PSL production appear to be preferentially directed towards cells which produce PSL; (5) cells which do not produce PSL are unable to successfully exploit cells which produce PSL. Taken together, this suggests that PSL is a social but relatively non-exploitable trait, and that growth within biofilms selects for PSL-producing strains, even when multiple strains can interact (low relatedness).

## IMPORTANCE

Many studies have shown that bacterial traits, such as siderophores and quorum sensing, are social in nature. This has led to a general assumption that secreted traits act as public goods, which are costly to produce and benefit both the producing cell and surrounding neighbours. Theory and experiments have shown that such traits are exploitable by asocial cheats, but we show here that this does not always hold true. We demonstrate that the *Pseudomonas aeruginosa* exopolysaccharide PSL provides social benefits to populations, but that it is non-exploitable, because most of the fitness benefits accrue to producing cells. More generally, our study highlights that experimental evidence is required to demonstrate the social nature of different bacterial traits.

## INTRODUCTION

The growth and proliferative success of many bacteria, including human pathogens, depend upon their ability to form biofilms in their respective environmental niches. Biofilms are multicellular three-dimensional structures, held together by extracellular matrix molecules that encapsulate cells and cause them to aggregate. These extracellular polysaccharides (EPS) which are secreted by the bacteria, typically function as adhesins that are used to attach cells to a surface and to maintain the three-dimensional biofilm structure, and sometimes aid in protection against a variety of stresses, including dehydration, antibiotics, and predators (1, 2). The production of EPS represents a problem from an evolutionary perspective (3), because it appears to be a type of co-operative behaviour that can potentially provide a benefit to all cells in the community (i.e. a ‘public good’), and not just to those that produce EPS. Consequently, the question arises: “what prevents the invasion of potential cheats that do not produce EPS?” Such cheats would presumably have a fitness advantage as they do not pay the metabolic cost of EPS production.

Two types of hypotheses have been suggested to explain why the costly production of EPS may be evolutionarily stable. The first hypothesis assumes that the production of EPS provides a benefit to the local population of cells, which can be exploited by cells not producing EPS (4, 5). This is directly analogous to a range of public goods that have been studied in bacteria, such as iron scavenging siderophore molecules and quorum sensing (6, 7). Both theory and experiments have shown that the production of exploitable public goods is favoured in spatially structured populations, which lead to a high relatedness between interacting cells, such that EPS producers tend to be aggregated and co-operating with other EPS producers (6–8). The second possibility, is that the production of EPS does not help biofilm growth *per se*, but it helps the EPS-producing lineage to outcompete other lineages that they are interacting with. For example, EPS producing cells might be able to spatially smother or displace non-producing lineages (9, 10).

From an evolutionary perspective, these two hypotheses are very different. The first involves an exploitable co-operative trait, whereas the second involves a trait that preferentially provides a benefit to other EPS producers. The different assumptions behind these hypotheses lead to different predictions. If EPS molecules are public goods, then monocultures of EPS producers will grow biofilms faster than non-producers, but the non-producers will be able to outcompete the producers in mixed biofilms (producers will have increases absolute growth, decrease relative growth) (4, 8). In contrast, if the function of EPS is to provide a competitive advantage, then we obtain the opposite prediction, that the non-producers will grow faster in mono-culture, and the EPS producers will grow faster in mixed biofilms (producers will have decreased absolute growth, but increased relative growth) (10). Support for these different hypotheses have been found examining different EPSs in different species – public good in *Pseudomonas fluorescens* and *Bacillus subtilis* (5, 11), and competitive advantage in *P. fluorescens, Pseudomonas aeruginosa* and *Vibrio cholerae* (12–15).

However, the generality of these explanations for the evolutionary stability of EPS production remains unclear. EPS molecules produced by different bacterial species can vary greatly in both their chemical structure and the biological roles they play within biofilms (16). EPS can even have consequences for other social traits, such as facilitating the transfer of other public goods (17). Furthermore, many species produce more than one type of EPS that are sometimes but not necessarily co-regulated. This means that there could be considerable variation in the social nature of different types of EPS, and for different bacterial species.

*P. aeruginosa* is an opportunistic pathogen that causes various biofilm infections such as chronic respiratory infections of cystic fibrosis (CF), keratitis, and chronic wound infections. It produces at least three different types of EPS molecules as major components of its biofilm matrix: alginate, PEL, and PSL polysaccharides (18, 19). Alginate production is inversely regulated with PSL (20, 21), and is not expressed to high levels in the majority of non-CF isolates (22, 23). In contrast, PSL is expressed in most *P. aeruginosa* natural and clinical isolates (22). PSL is a crucial adhesive scaffolding component of the biofilm matrix, promoting both cell-to-cell interactions and surface attachment (24–26). PSL also has a unique function as an intercellular signalling molecule (27), underscoring its roles in social evolutionary interactions. Here we test the social nature of PSL and find that it is a social trait, which provides benefits at the individual and group level, but that it cannot be successfully cheated by individuals who do not produce it. Our results therefore point to a scenario that is distinct from either of the hypotheses that have previously been proposed. A broad implication of our work is that not all components of the biofilm matrix act as shared resources.

## RESULTS

### PSL provides a population level benefit in biofilms

We first tested whether PSL provides fitness benefits to a *P. aeruginosa* population growing in biofilms or non-biofilms compared to that of populations of mutants that do not produce PSL. To this end, we measured the amount of biofilm and non-biofilm biomass produced over 4 days’ growth for PSL producer and non-producer strains. In order to simultaneously monitor both unattached and biofilm sub-populations within a microcosm, we modified a previously described bead method (28). This model allows us to grow biofilms on 7 mm plastic beads in test tubes, harvest biofilm cells from the beads, and directly aspirate cells that were not attached to beads from the liquid media of the same culture (Fig. S1). Cells not attached to beads likely include all or combinations of any of the following: unattached multicellular aggregates, free-swimming ‘planktonic cells’, cells defective in biofilm formation, and cells dispersed from biofilms. Since these cell types can all be transient and inseparable or indistinguishable, in this paper we unify all of these types of cells and refer to this sub-population as ‘unattached cells’. We used two strains: a PSL+ strain that constitutively produces PSL, and a PSL-mutant that produces no PSL. PSL expression has been shown to induce an increase of the intracellular concentration of the secondary messenger molecule c-di-GMP, which controls multiple biofilm-associated genes (27). Consequently, to ensure that our results are due to PSL production specifically and not downstream c-di-GMP-dependent pleiotropic effects, we constitutively elevated c-di-GMP (Δ*wspF* background) in both our strains, but nevertheless ultimately found that our results are c-di-GMP-independent because qualitatively identical results were recorded using non-Δ*wspF* backgrounds (Fig. S2). Δ*wspF* strains constitutively upregulate both *psl* and *pel* transcription (29), thus maximising the phenotypic effects of the EPS in question. It is noteworthy that Δ*wspF* mutants are frequently selected for in *in vitro* biofilms and *in vivo* biofilm-related infections (30, 31). Furthermore, to ensure our results were solely dependent on PSL, we mutated the other c-di-GMP co-regulated EPS gene locus *pel*. Thus, henceforth in this paper, unless specified otherwise, our “PSL+ strain” is Δ*wspF* Δ*pel*, and the “PSL-strain” is Δ*wspF* Δ*pel* Δ*psl*. Later, when we address the PEL polysaccharides, the “PEL+ strain” is Δ*wspF* Δ*psl*, and the “PEL-strain” is Δ*wspF* Δ*pel* Δ*psl*.

Our results show that PSL provides a population level benefit in biofilms but not in unattached populations (Fig. 1A). Consistent with previous reports (22, 24–26, 32), we found that over 4 days of growth, PSL mutants formed significantly less biomass and therefore poorer biofilms compared to the corresponding PSL+ strain. In contrast, we found no significant differences in the final population density between the PSL+ and the PSL-strains in the unattached population of cells.

**Fig. 1.**
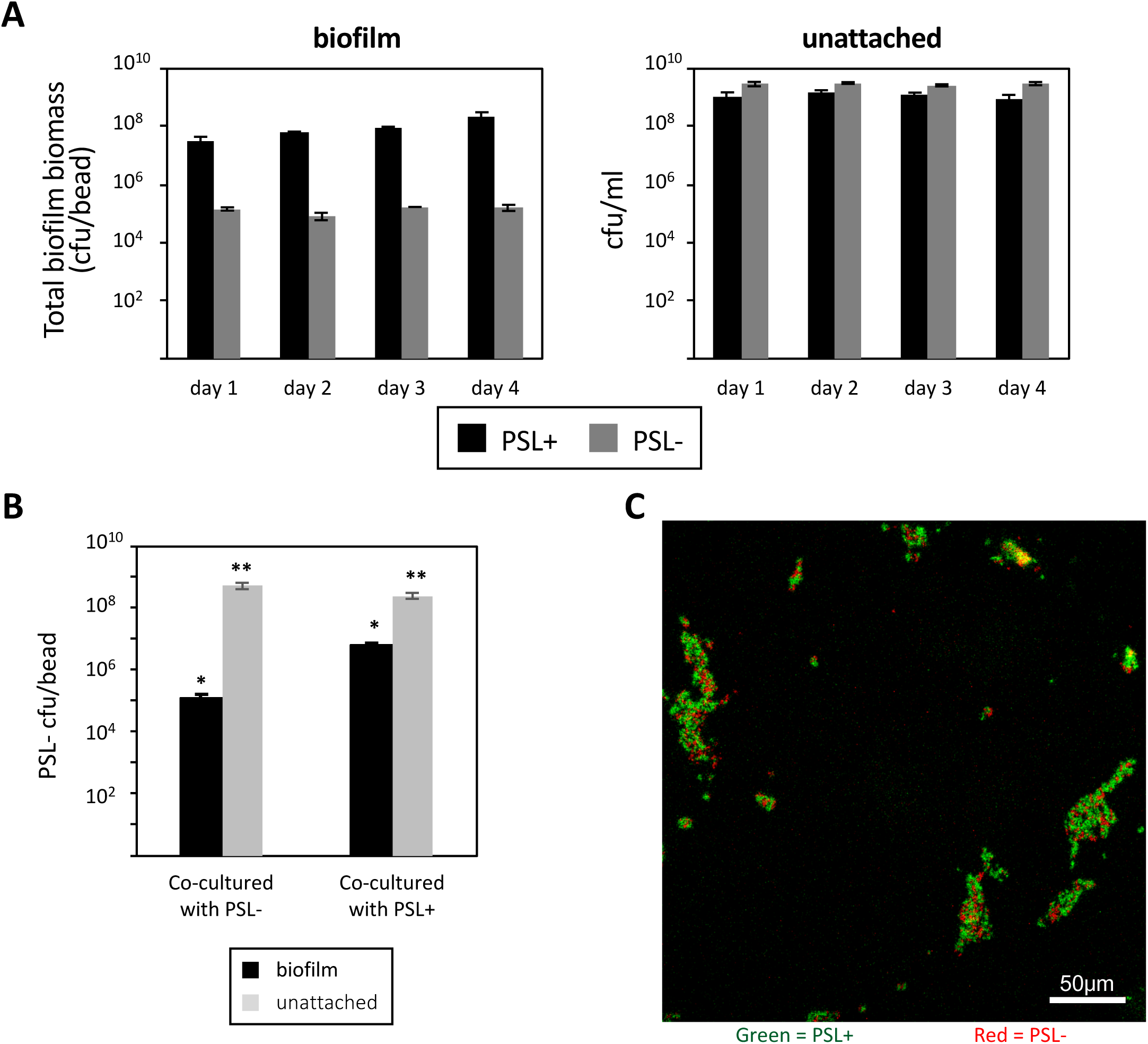
PSL production provides community benefits to cells grown in biofilms. (*A*) PSL-strain is significantly defective in biofilm formation on beads compared to PSL+ (*F*_(1,15)_ = 11.26, *p* = 0.0043, n = 3), but no major differences are seen between their growth in the unattached fractions (*F*_(1,16)_ = 0.67, *p* = 0.4251, n = 3). (*B*) PSL-co-cultured with PSL+ at a 1:1 ratio increases their proportional numbers in a biofilm (* *p* < 0.0001; ** *p* > 0.08; n=4). (*C*) Confocal micrograph image of surface attached populations of PSL-/PSL+ 1:1 co-cultures. PSL-cells (red) co-aggregate with PSL+ cells (green).

In order to confirm that our unattached population results were not biased by cells that had detached from biofilms, we also performed an experiment where we added no beads and therefore the analyses were all performed on purely unattached cells. Again we found no significant differences in the final population density between PSL+ and PSL-strains (Fig. S3). Since we found that a significant biomass was achieved after 1 day of biofilm growth, and in order to minimise the occurrence of rapid spontaneous mutations that can develop and accumulate in mature biofilms of *P. aeruginosa* (32), we carried out all subsequent experiments with 24 hour cultures.

### PSL provides social benefits in biofilms

We then asked whether the production of PSL provides social benefits to other cells within a biofilm. We tested this in unattached populations and in biofilms, by growing the PSL-strain with either other PSL-cells or PSL+ cells in a 1:1 ratio. We found that approximately 100-fold more PSL-cells could attach and form biofilms in the presence of PSL+ compared to when PSL-was co-cultured with PSL- (Fig. 1B). In contrast, in unattached populations, we found that the fitness of PSL-cells was not influenced by co-culturing with PSL+ (Fig. 1B). This indicates that, in biofilms, the production of PSL provides some benefits to cells that do not produce PSL, but not in unattached populations. Consistent with PSL+ providing a benefit to PSL-cells, we found that when we co-cultured PSL+ and PSL-strains, PSL-cells co-aggregated with PSL+ cells and incorporated themselves into the biofilm during the early stages of biofilm formation (Fig. 1C).

### PSL mutants do not act as social cheats within biofilms

We next tested whether PSL-strains could act as social cheats (33), i.e. whether these strains increased in frequency when growing in mixed cultures with the PSL+ strain. We varied the starting ratios of PSL-:PSL+ cells from 0.1 to 10,000, because theory predicts that the fitness of cheats should be frequency-dependent, with cheats being better able to exploit co-operators when the cheats are rarer (34). In biofilms, we found that the relative fitness of the PSL-cells was either equal to or lower than that of the PSL+ strain, suggesting that the PSL-cells were not able to outcompete the PSL+ cells (i.e. to cheat). The relative fitness of the PSL-strain was negatively correlated with its starting frequency in the population (Fig. 2A). At high starting frequencies of PSL+, the relative fitness of the PSL-strain was not significantly different from that of the PSL+ strain. As the starting frequency of the PSL-strain was increased, the relative fitness of the PSL-strain became lower than that of the PSL+ strain. Thus, the PSL-strain does not outcompete the PSL+ strain at any starting frequency, and at high starting frequency it is in fact outcompeted by the PSL+ strain. In contrast, we found that in unattached populations, the relative fitness of the PSL-strain did not differ significantly from the PSL+ strain, irrespective of the starting ratio (Fig. 2A).

**Fig. 2.**
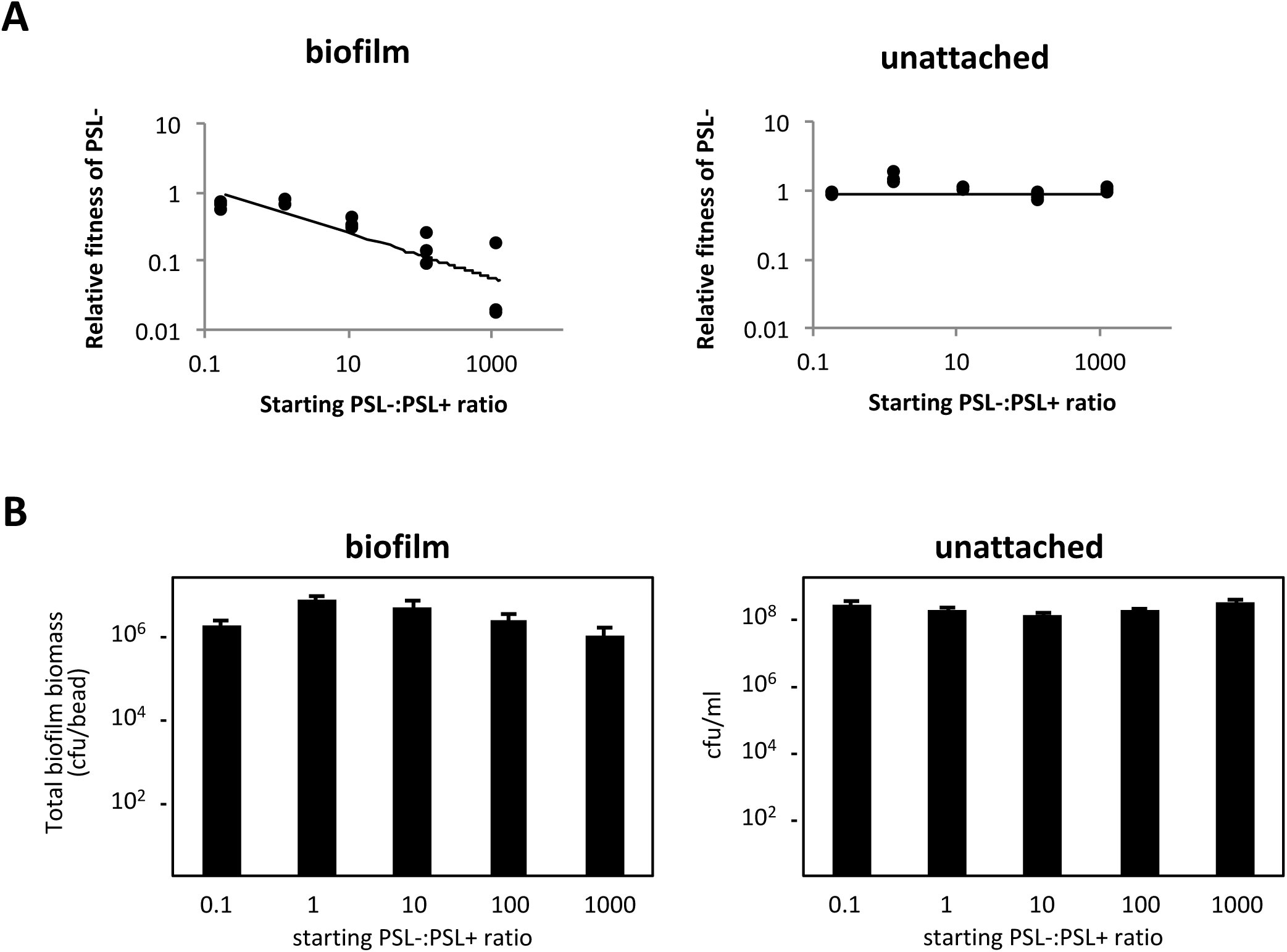
PSL mutants are not social cheats within biofilms. (*A*) The relative fitness of PSL-cells was equal or lower than the PSL+ strain across all starting frequencies of the mutant in mixed cultures in biofilms (*t*(13) = −3.242, *p* = 0.0064). PSL-strain fitness was equal to PSL+ strain fitness across all starting frequencies in unattached populations (*t*(13) = −0.741, *p* = 0.4716). (*B*) PSL-/PSL+ co-culture biomass remained similar across all starting ratios in both biofilm (*t*(13) = −1.769, *p* = 0.1004) and unattached populations (*t*(13) = 1.881, *p* = 0.0825).

We also tested whether the presence of the PSL-strain adversely affected biofilm production in mixed cultures. Given that the PSL-strain is unable to form biofilms to the same biomass level as the PSL+ strain (Fig. 1), we expected that a high starting frequency of the PSL-strain would adversely affect biofilm production. In contrast to this expectation, we found that the final biomass did not significantly change as we varied the starting ratio of PSL-:PSL+ cells from 0.1 to 1000 (Fig. 2B). This result indicates the final biomass is determined by the PSL+ cells, which dominate long-time biofilm growth independently of the starting ratio between PSL-and PSL+. To allow us to microscopically visualise growth of the PSL-and PSL+ strain in co-inoculated biofilms, we performed a biofilm flow cell experiment with a mixed PSL+ and PSL-population. Consistent with Fig. 1C, PSL-cells appeared to co-aggregate with PSL+ cells and incorporate themselves into the early-stage biofilms (Figs. 3A and B). However, over time, the PSL+ cells covered and outcompeted the PSL-cells (Figs. 3C and D).

**Fig. 3.**
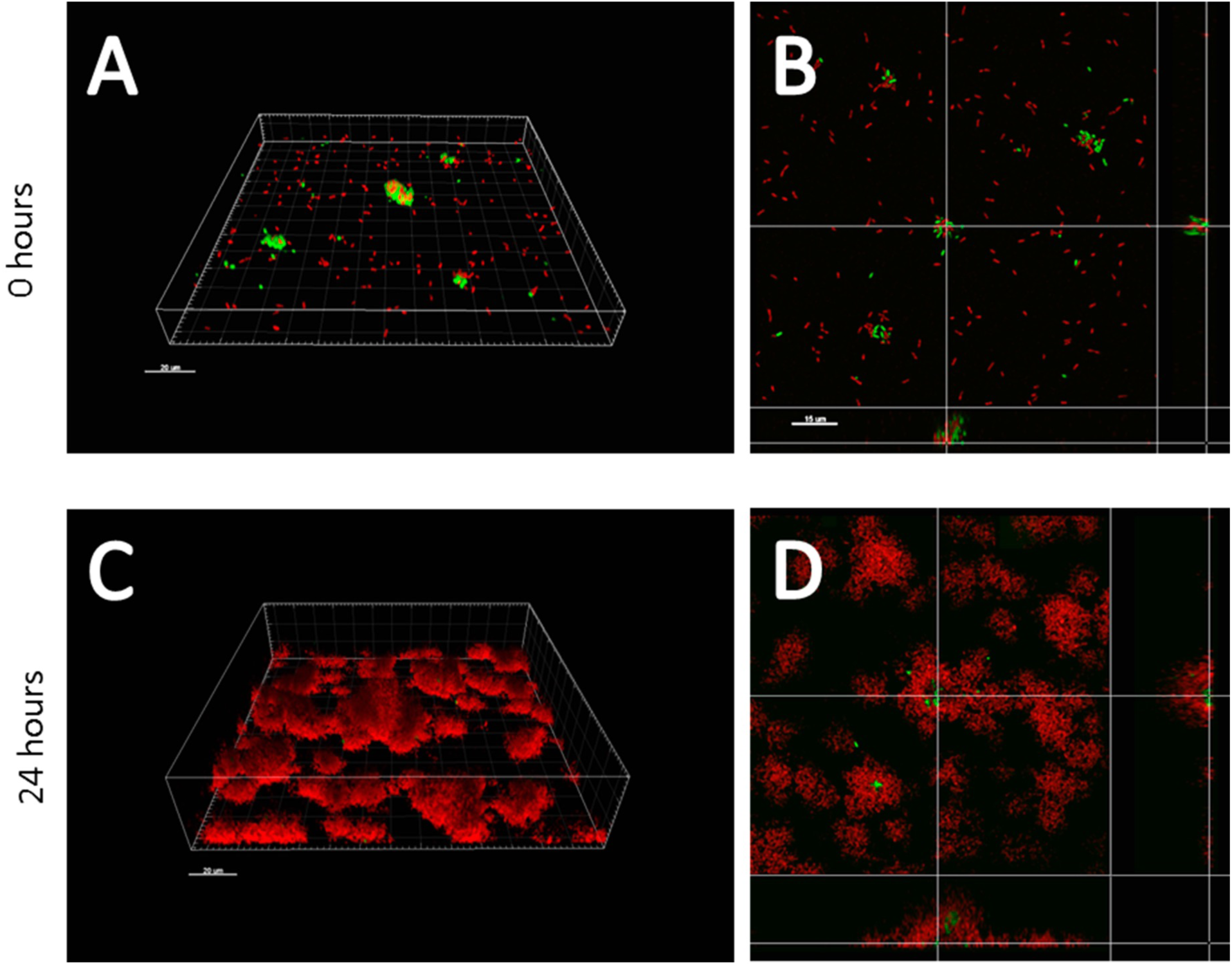
The PSL+ strain smothers and outcompetes the PSL-strain during biofilm growth. Confocal micrograph image of PSL-/PSL+ (1:1) mixed population biofilms at 0 hours (*A* and *B*) and 48 hours (*C* and *D*). PSL-cells are only attached to the surface as co-aggregates with PSL+, and they are eventually outcompeted by PSL+ when the biofilm matures. *A* and *C* are 3D rendered images and *B* and *D* represent corresponding open-box top-down views. Red = PSL+ cells. Green = PSL-cells.

### PSL provides population level benefits against antibiotics

PSL is known to play a role in tolerance to antibiotics including aminoglycosides (32, 35) (Fig. S4). We tested the social consequences of this tolerance by examining the effect of adding gentamicin at a concentration previously known to affect biofilm (100 μg/ml) (36) to populations that contained a variable ratio of PSL-mutants. In the absence of gentamicin, we found that the frequency of PSL-mutants did not influence the biomass of biofilm, consistent with Fig. 2B (Fig. 4A). In contrast, when we added gentamicin, the biofilm biomass was negatively correlated with the fraction of PSL-mutants in the starting population (Fig. 4A). When there were more PSL-than PSL+ cells in the starting population, the addition of gentamicin led to a significant drop in viable biofilm biomass. In cases where the starting ratio of PSL-:PSL+ was greater than 10, the addition of gentamicin reduced the population to below our threshold detection level. Examining in more detail the populations which showed partial killing, we found that PSL+ cells had a significantly higher level of survival than the PSL-cells (Fig. 4B).

**Fig. 4.**
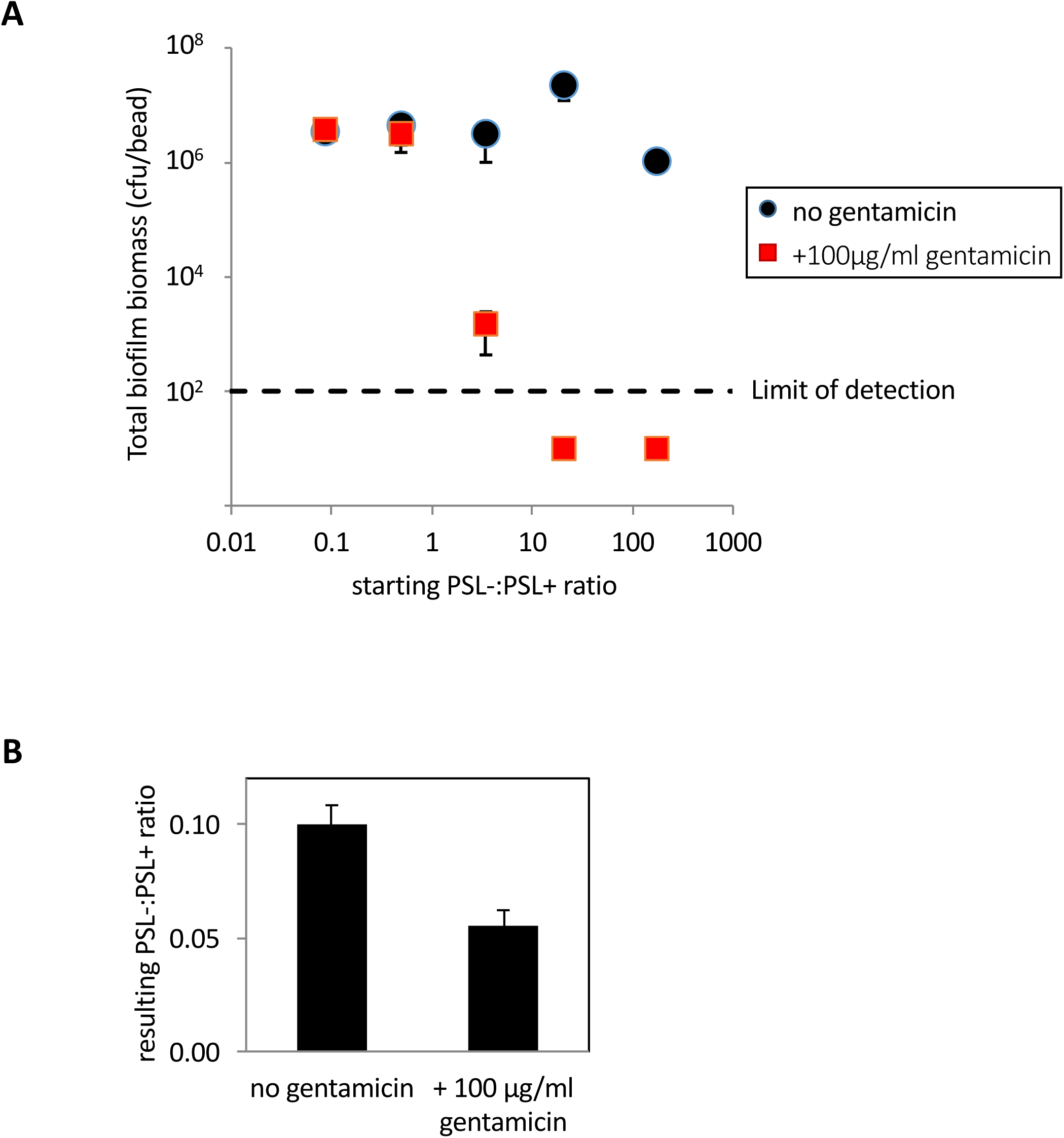
PSL production results in increased tolerance towards antibiotics. (*A*) PSL-/PSL+ mixed biofilms yielded consistent biomass regardless of the starting ratio of the two strains (*t*(13) = −0.732w, *p* = 0.4769, n=3), treatment of the mixed biofilm with a high concentration of gentamicin eliminated the entire biofilm population when the starting proportion of PSL-cells increased above PSL+. (*B*) In the intermediate PSL-:PSL+ co-culture ratio, where mixed biofilm was partially eradicated, there were less PSL-cells in the surviving biofilm in the presence of gentamicin (*t(*4) = 4.07, *p* < 0.02, n = 3).

### Relatedness does not influence selection for PSL

We complemented the above fitness assays with a multi-generational selection experiment. A number of previous studies on microbial social traits such as siderophore production and quorum sensing, have shown that the relative fitness of individuals that do and do not perform social traits depends upon population structure (6, 7, 37, 38). Specifically, structured populations, with a relatively high relatedness between cells (low strain diversity), favour genotypes that support co-operative traits. Conversely, populations with relatively low relatedness between cells, (high strain diversity) favour less co-operative cheating genotypes. These results are as predicted by social evolution theory (39, 40). In contrast, our observation that PSL-is not able to cheat PSL+ in mixed cultures, suggests that relatedness might not matter for the evolution of PSL in the same way.

We tested this hypothesis with a selection experiment over 5 days, in which we started with a mixed population of PSL+ and PSL-cells, and manipulated both relatedness (low/high) and whether cells were growing in biofilms (on beads) or in cultures with no beads (the entire population is unattached) (Fig. S5). In each round of growth, we subdivided the population into a number of subpopulations. We varied relatedness by initiating each subpopulation with either a single clone, to give relatively high relatedness, or with multiple clones, to give relatively low relatedness. Consequently, in the high relatedness treatment the PSL+ and PSL-strains become segregated into separate populations, while the low relatedness treatment keeps them mixed together in the same population. We started our experiments with a 1:1 ratio of PSL-:PSL+, and carried out six rounds of growth.

We found that, in contrast to previous selection experiments on microbial social traits, relatedness had no influence on the outcome of our selection experiment (Fig. 5). Instead, we found that the relative fitness of the PSL-and PSL+ strains were determined by whether the sub-populations were grown as biofilms or in unattached populations. Specifically, the PSL+ strain was significantly favoured in biofilms but not in unattached populations in bead-less cultures (Fig. 5). Consequently, our selection experiment provides further support for our conclusion that PSL+ is favoured in biofilms, and that PSL-strains are unable to cheat PSL+ strains during growth in biofilms.

**Fig. 5.**
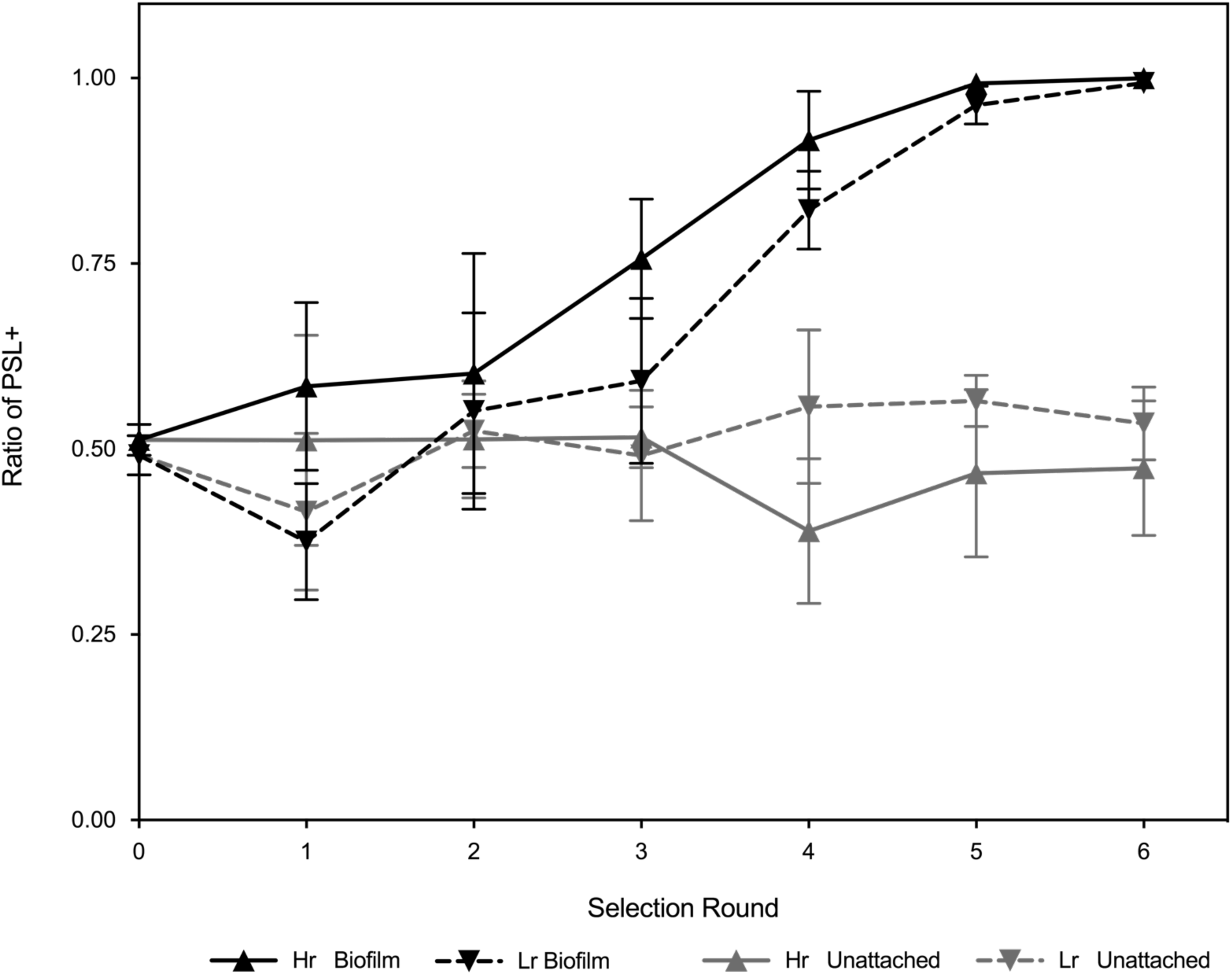
PSL production is favoured in high and low relatedness conditions in biofilms. PSL+ cells have a selective advantage under conditions promoting biofilm formation (black) (*F* _[1,20]_ = 542.7, *p* < 0.0001, n=6), however this observation is not reciprocated for unattached conditions (grey). The concurrent loss of PSL-cells from both low related and high related biofilm populations through subsequent selection rounds, suggests there is a strong selection towards PSL+ strains (under biofilm-promoting conditions only) as PSL- strains are unable to cheat on PSL+ strains.

## DISCUSSION

We examined the social nature of the PSL polysaccharide produced by *P. aeruginosa* and found that: (1) PSL production provides a benefit at the group level to cells growing in biofilms (Fig. 1A); (2) PSL production provides no benefit or cost to cells growing in unattached populations (Figs. 1A and 2A); (3) PSL-strains are able to grow better as biofilms when in a mixed culture with PSL+ cells (Fig. 1B); (4) PSL+ strains are able to outcompete PSL-strains when in mixed culture biofilms, such that PSL-strains cannot successfully exploit (cheat) the PSL production of the PSL+ strain (Fig. 2A); (5) the relative fitness of the PSL-strain is lower when the PSL+ strain is rarer (negative frequency dependence; Fig. 2A); (6) the presence of PSL-cells has a negligible influence on overall biofilm productivity (Fig. 2B); (7) biofilms containing a higher proportion of PSL+ cells are less susceptible to antibiotics (Fig. 4A); (8) PSL+ cells are better able to survive antibiotics when growing in a mixed culture biofilm with PSL-cells (Fig. 4B); (9) in a selection experiment, the relative success of PSL+ versus PSL-was not influenced by relatedness, for growth in biofilms (Fig. 5).

Overall, our results suggest that in biofilms, the production of PSL is a social trait, which provides benefits at the individual and group level, but that it cannot be successfully cheated by individuals who do not produce it. This is despite the fact that the PSL-strain is able to gain some benefit from the presence of the PSL producing strain in the biofilm matrix (Fig. 1). Our experiments suggest that the fitness benefits of producing PSL accrue mainly to the producing cell and/or to other PSL+ cells. This contrasts with previous studies on social traits in *P. aeruginosa*, where it has been shown that siderophore production and quorum sensing (QS) are social traits that can be readily exploited by cheats in *in vitro, in vivo*, and in biofilm experimental systems (6, 7, 38, 41). We suggest that this is because siderophores and most QS-dependent public goods are relatively freely diffusible secreted products that provide shared benefits to both producing and non-producing cells. In contrast, the primary function of PSL is adherence to surfaces and to other cells, PSL likely has limited diffusive properties in the biofilm biomass. PSL was previously shown to localise primarily to the periphery of the biofilm microcolonies, encapsulating the cells (42). In addition, some PSL is known to be tightly associated to the cell surface (43), although some is likely to be diffusible, and serves as an intercellular signalling molecule (27). This signalling property of PSL also promotes co-ordinated, community-level gene expression towards increased production of PSL and other biofilm-associated factors by driving its own expression in a feed-forward manner, similar to that of an auto-induction regulatory circuit.

Our results do not match either of the hypotheses which have been suggested to explain the evolutionary stability of EPS. Consistent with the public good hypothesis, we found that PSL production led to higher growth in monocultures, and that it provided a social benefit (Figs 1A, 1B & 4A) (4, 5). However, in contrast to what we would expect with a public good, we found that: (a) PSL-strains could not successfully exploit (cheat) the PSL production of the PSL+ strain (Fig. 2A); (b) the presence of PSL-cells had a negligible influence on overall biofilm productivity (Fig. 2B); and (c) the relative success of PSL+ versus PSL-was not influenced by the relatedness between cells within biofilms (Fig. 5). Consistent with the competitive advantage hypothesis, we found that PSL+ strains are able to outcompete PSL-strains when in mixed culture biofilms (Fig. 2A) (10).2A). However, contradictory to the assumptions and predictions of the competitive advantage hypothesis, we found that: (a) PSL provides an advantage, not a disadvantage in biofilm growth, with PSL production providing a benefit at the group level to cells growing in biofilms (Fig. 1A); (b) PSL production provides a benefit to non-producers, with PSL-strains able to grow better as biofilms when in a mixed culture with PSL+ cells (Fig. 1B); (c) the benefit of producing PSL was increased by the presence of competitors (Fig. 4A).

Consequently, PSL appears to align with a different type of social trait, sharing some features of both hypotheses previously suggested. PSL is closer to a co-operative public good, as it provides a benefit at the group level, leading to faster growing and better defended biofilms, and this benefit can be exploited by non-producers (Figs. 1 & 4). The extent, however, to which it is exploitable by non-producers is very small, such that PSL+ outcompete PSL-in mixed biofilms (Figs. 2, 3 & 5). This outcompeting in mixed biofilm result is that predicted by competitive advantage models (10), but it arises for a different reason: the inability of non-producers to benefit from a co-operative trait that increases absolute growth, rather than a trait that reduces absolute biofilm growth, but provides a relative competitive advantage (10).. It is hard to disentangle whether the ability of PSL+ to outcompete PSL-in mixed biofilms is a property of the oxygen/nutrient gradient, or whether it is the property of differing adhesive strengths between PSL+ and PSL-cells, as PSL-cells are less adhesive and are more prone to be detached from the biofilm biomass in constant flow conditions when exposed on the surface. A greater adhesive ability can allow strains to displace competitors and gain an advantage in biofilms (9), as evidenced by more evenly spaced existence of PSL-cells amongst PSL+ aggregates within an early developing biofilm under no or low flow conditions (Fig. 1C).

Our experiments suggest that PSL production cannot be cheated, and hence should be favoured even in low relatedness mixed cultures. We tested the effect of relatedness on selection for PSL production in biofilms, using a selection experiment (Fig. 5). In our experiment, high relatedness implied that cultures were established with either PSL+ or PSL-strains. Under these conditions, cells will interact with genetically identical cells, and PSL-cells cannot interact with PSL+ cells. In contrast, low relatedness implies that cultures can be comprised of both PSL+ and PSL-strains, such that PSL-cells can theoretically exploit the PSL+ cells. We found that relatedness had no effect on our results. PSL+ strains were favoured in conditions of high and low relatedness in biofilms but not in non-biofilm populations. Therefore in mixed culture biofilms, a PSL producer always wins, in contrast to previous work on other social traits, such as siderophore production and QS (6, 7, 38, 41). In these previous studies, relatedness matters, because at low relatedness, the uncooperative cells are able to interact with the co-operative cells, allowing the uncooperative strains to exploit (cheat) the co-operative wild type cells (44). Our selection experiment provides further evidence that the benefits of producing PSL directly affect the producing cells and that PSL production is not exploitable by non-producing cheats.

We further demonstrated that PSL helps protect biofilm communities against environmental challenge. When biofilms contain a high proportion of PSL-cells, this leads to a significant increase in the susceptibility to antibiotics (Fig. 4A). Within these biofilms, PSL+ cells showed a higher level of survival than the PSL-cells (Fig. 4B), indicating that the protection provided to the cells by PSL is most beneficial for the producing cells. This is likely due to PSL being both cell-associated and released into the media (20). While PSL-cells can readily access released PSL, they do not have access to cell-associated PSL. One possibility is that there are functional differences for providing antibiotic tolerance between cell-associated PSL and released PSL. Alternatively, the relative PSL concentration may simply be higher in closer proximity to the cell surface, providing a higher level of protection against antibiotics. Shared benefit appears to be provided to the community as a whole including non-producing cells when they are rarer. This may be because biofilm biomass appears to be entirely encapsulated with possibly the released form of PSL (42). The exact mechanism of antibiotic tolerance by PSL is not known, and this therefore remains an open question.

Arguments have been made previously, that *P. aeruginosa* cells, even when they are in the ‘planktonic’ phase, have biofilm-like features (45). In our bead biofilm model, the unattached cell population, particularly those cells which constitutively express EPS, contain large numbers of cellular aggregates, as seen previously in shaken liquid cultures (27, 42). However, our results clearly distinguish the social evolutionary differences between the cells in the biofilm and unattached cells, thus providing evidence that unattached cells, whether they are aggregated or not, are definitively distinct from surface-attached biofilm cells. This raises the question whether aggregates represent a type of biofilm, are ‘biofilm-like’, or represent a different growth mode altogether, and our results are strongly suggestive for the latter. This is an important question since *P. aeruginosa* biofilms growing on infected contact lens, catheters, and other medical implants form surface-associated biofilms, but within the cystic fibrosis sputum, the bacteria are thought to grow as suspended aggregates (46–48).

Most strains of *P. aeruginosa*, like many other microbial species, are capable of producing multiple biofilm EPS molecules (22, 49). Of these, PSL and PEL are co-regulated by several intracellular regulatory factors including c-di-GMP (50) and both contribute to biofilm development in a majority of *P. aeruginosa* strains including PAO1 (22). Intriguingly, in additional experiments, we found that the social aspects of PEL production are different from those of PSL, pointing towards PEL as being a completely private and non-social good (Fig. S6). This is in contrast to recent work inspecting the competitive fitness of PEL producers using the PA14 strain, which showed community-level benefits in some circumstances (13). PA14 has a deletion mutation from the *psl* promoter through the *pslD* gene, and the nature of biofilm formation in this strain, which uses PEL polysaccharide as the sole EPS, appears to have diverged from that of other *P. aeruginosa* isolates. It is possible that PA14 has somehow evolved PEL to compensate for the loss of PSL by altering not only its expression patterns (49), but also its social evolutionary roles. These results highlight the importance of recognising diverse roles and effects of different EPS as social traits within biofilms. The extracellular matrix is the material that keeps microbial cells together in a biofilm community, influencing their social interactions and evolution. As such we believe it holds an important foundation for our understanding of the multicellularity of unicellular organisms.

## MATERIALS AND METHODS

### Bacterial strains and growth conditions

The bacterial strains and plasmids we used and constructed for this study are listed in Table S1. We propagated *Escherichia coli* and *P. aeruginosa* strains in Lysogeny Broth (LB) at 37°C unless otherwise specified. The concentrations of antibiotics we used for *E. coli* were: 50 μg/ml carbenicillin and 10 μg/ml gentamicin, and for *P. aeruginosa:* 300 μg/ml carbenicillin and 100 μg/ml gentamicin. We induced P_BAD_-*psl* strains for PSL over-expression by adding 1 % w/v L-arabinose. PSL is involved in a feed-forward regulation system where the expression of PSL induces the intracellular levels of c-di-GMP, which in turns over-expresses PSL (27). To avoid c-di-GMP being inadvertently triggered in our biofilm system, which would potentially affect the interpretations of our results when other c-di-GMP-induced gene products become involved, all strains used in our studies were conducted using a Δ*wspF* mutation background. Δ*wspF* mutants express biologically maximal intracellular c-di-GMP levels due to the constitutive activation of diguanylate cyclase WspR (29).

### Fluorescent strain construction

We chromosomally labelled *P. aeruginosa* strains with either GFP or mCherry using the Tn7 delivery system (51). We electroporated 1 μl pTNS3 + 1 μl pUC18-mini-Tn7T2-PA1/04/03::*gfp* or pUC18-mini-Tn7T2-PA1/04/03::*mcherry* into a respective electrocompetent *P. aeruginosa* strain as previously described (52) and selected for gentamicin resistance. We removed the construct backbone by electroporating 1 μl pFLP2 and selecting for carbenicillin resistance. Upon sucrose counter-selection to identify the loss of the *sacB* gene-containing pFLP2 by streaking for single colonies on LB (no salt) plus 10 % sucrose at 30°C, we confirmed the strains for carbenicillin and gentamicin sensitivities.

### Bead biofilm culture system

We grew biofilm cultures on plastic beads as previously described (28) but with several modifications (Fig. S1). We used this system because it allowed us to simultaneously study biofilm and unattached populations. This system differs from traditional flow cell models in that it is closed, and unattached populations are not lost due to flow. Therefore any differences that we observe are reflective of biofilm vs unattached populations instead of ‘biofilm cells being the only ones left;’, which can occur in a flow cell system. We streaked *P. aeruginosa* strains to obtain single colonies on M9 + 3.062 g/l sodium citrate (M9 citrate) agar at 37°C for 2 days. We picked individual fresh colonies and grew these to logarithmic phase (OD600 ≈ 0.5) in M9 citrate broth at 37°C with shaking. We then diluted these cultures to OD600 ≈ 0.05 into 3 ml M9 citrate containing one or more (up to three) 7mm plastic beads (Lascells) in standard culture tubes into independent triplicates. We then grew the cultures at 37°C with shaking at 200 rpm for 24 hours. We directly aspirated unattached cells from the liquid broth portion of the culture. We collected biofilm cells by retrieving the beads from the tubes, and gently washed the beads in 10 ml PBS by five inversions which we repeated three times. We determined the colony forming units (CFU) of the biofilms by recovering the cells by water bath sonication in 1 ml PBS for 10 min. We vortexed the sonicated samples for 10 sec prior to serial dilution and plating.

### Antibiotic killing assays

For antibiotic killing assays (Fig. 4), we first grew bead biofilm cultures for 24 hours in the absence of gentamicin. Half the cultures were then introduced to 100 μg/ml gentamicin (final concentration), and the other half were left untreated. We then further incubated the cultures for 24 more hours at 37°C with shaking prior to harvesting the cells for analyses.

### Determining ratios of PSL+ and PSL-cells in mixed cultures

To distinguish between PSL+ and PSL-cells, we introduced a single chromosomal copy of constitutively expressed GFP or mCherry genes to each strain. We then used quantitative real-time PCR and primers specifically designed against GFP or mCherry genes to accurately assess the numbers of each strains present in a sample. In order to avoid the effects of PEL, which is co-regulated with PSL, interfering with or convoluting PSL-dependent phenotypes, we conducted all experiments using Δ*pel* mutant backgrounds.

### Quantitative real-time PCR

We collected genomic DNA (gDNA) from biofilm cells from washed beads without the sonication steps by directly applying lysing reagents to the beads. We isolated unattached bacterial gDNA by pelleting 1 ml of the liquid cultures. We then stored pellets and beads at −20°C until DNA extraction. We extracted gDNA using a GenElute^TM^ Bacterial Genomic DNA Kit (Sigma) according to the manufacturer’s protocol except for the elution buffer which we diluted 100-fold to prevent EDTA interfering with the subsequent DNA polymerase reactions. We performed real-time PCR as previously described (27) using Syber Green PCR Master Mix (Applied Biosystems). The primers we used are listed in Table S2. We generated standard curves for *gfp* and *mcherry* genes using quantitatively determined gDNA from Δ*pel* Δ*psl* GFP strain and Δ*pel* Δ*psl* mCherry strain respectively.

### Biofilms grown on coverslips and confocal microscopy of biofilms

Bacteria were cultured as described in the bead biofilm system, but instead of inoculating into bead-containing tubes, cultures were diluted into 1 ml M9 citrate in uncoated polystyrene 24-well plates. Each wells contained one sterile 10 mm diameter glass coverslip. Biofilms were grown in 37°C with 200 rpm shaking for 24 hours. The glass coverslips were removed by forceps, gently washed three times in 10 ml PBS, and immediately mounted on a glass slide for confocal microscopy by Zeiss LSM 700. Images were acquired and analysed using Zeiss’s ZEN software.

### Flow cell growth and confocal microscopy of biofilms

We grew mixed Δ*pel* Δ*psl* and Δ*pel* aggregates in a continuous flow cell system as described previously (53). We produced the mixed aggregates by mixing exponential growing liquid cultures, adjusted to an OD_600_ of 0.001, 1:1 in LB media. This mix was then incubated overnight shaken at 180 rpm. We diluted the aggregate-containing culture to an OD600 of 0.01 before inoculating the flow cells with 27G syringes. We used a flow rate of 3 ml/hour of M9 minimal media buffered with 10 % v/v A10 phosphate buffer supplemented with 0.3 mM glucose. With a Zeiss Imager.Z2 microscope with LSM 710 CLSM and the accompanying software Zeiss Zen 2010 v. 6.0, we found aggregates containing a mix of the GFP-and the mCherry-tagged cells. We imaged these aggregates over time with z-stack intervals of 1 μm. 3D projections of images were produced in Imaris (Bitplane, Switzerland). We quantified biomass of the two tagged populations with the open source software FIJI (LOCI) and the plugin Voxel_Counter (NIH).

### Biofilm selection experiment

In order to determine potential fitness of PSL production, we performed a selection experiment over 6 rounds of growth, with four different treatment regimens; relatively low and relatively high relatedness (Lr and Hr, respectively) (Fig. S5). The Hr treatment started with three 3 ml subpopulations of diluted logarithmic phase PSL+ cells and PSL-cells (n = 6). In contrast, the Lr treatment started with six subpopulations of a 3 ml 1:1 mix of PSL+ and PSL-cells (n = 6). We grew cultures at 37°C with shaking at 200 rpm for 24 hours in the presence/absence of plastic beads. We recovered biofilm cells as detailed previously and pooled together prior to serial dilution and plating. We recovered 1ml of unattached cells from the liquid media *in lieu* of plastic beads, and subjected to the same treatment (omitting the three wash stages). We determined the ratio of PSL+ and PSL-cells in each treatment regimen phenotypically; PSL+ and PSL-cells have rugose and smooth colony morphologies, respectively. For Hr treatments we then randomly selected six clones to inoculate 3 ml M9 citrate and progress to the next round, whereas for LR treatments, we randomly took a sweep of multiple colonies. We grew cultures at 37°C with shaking at 200 rpm for 24 hours in the presence/absence of plastic beads adding an additional round of selection. We repeated this experiment six times, with six selection lines per treatment regimen.

### Mathematical formula and statistics

We determined relative fitness (*w*) of the PSL-strain using the following formula: w = x_2_ (1 − x_1_)/x_1_ (1 − x_2_), where x_1_ is the starting mutant proportion of the population and x2 is the end mutant proportion (34). All error bars denote standard errors of the mean. All statistical tests including unpaired *t* tests, linear regression analyses and one-way and two-way analyses of variants were performed on VassarStats (http://vassarstats.net/).

## ACKNOWLEDGEMENTS

The authors would like to thank Boo Shan Tseng and Matt Parsek for their generous gifts of GFP and mCherry chromosomal labelling constructs. Y.I. would like to thank Dr Andrew Preston for the MC sponsorship and scientific support. This work was supported by a Human Frontier Science Program Young Investigators grant to V.G., R.J.A., T.B. and S.P.D. (RGY0081/2012), a NERC grant to S.P.D. and S.A.W. (NE/J007064/1), a European Research Grant to S.A.W., an FP7 Marie Curie Fellowship grant to Y.I. (PIIF-GA-2012-329832), a grant from the Lundbeck Foundation to T.B. and K.N.K. and Royal Society University Research Fellowships to S.P.D. and R.J.A.

**Author contributions.** Y.I., A.E.L.R. and K.N.K. performed research; Y.I., A.E.L.R., K.N.K., T.B., R.A., G.M., V.D.G., J.H., S.A.W. and S.P.D. designed the research and analysed the data; Y.I., S.A.W. and S.P.D. wrote the manuscript.

**Competing interests.** The authors declare no competing or conflicts of interest associated with this manuscript.

---

**Fig. S1.**
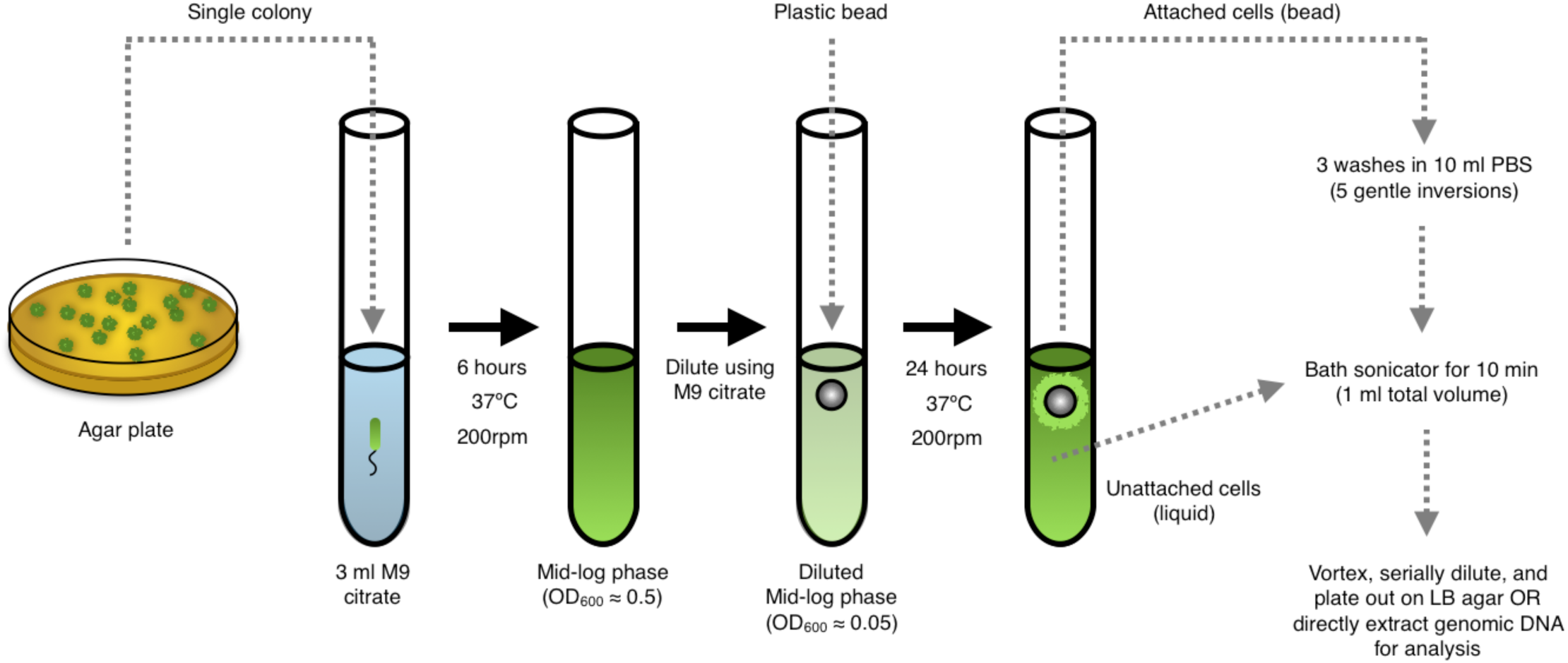
Schematic of the bead biofilm system.

**Fig. S2.**
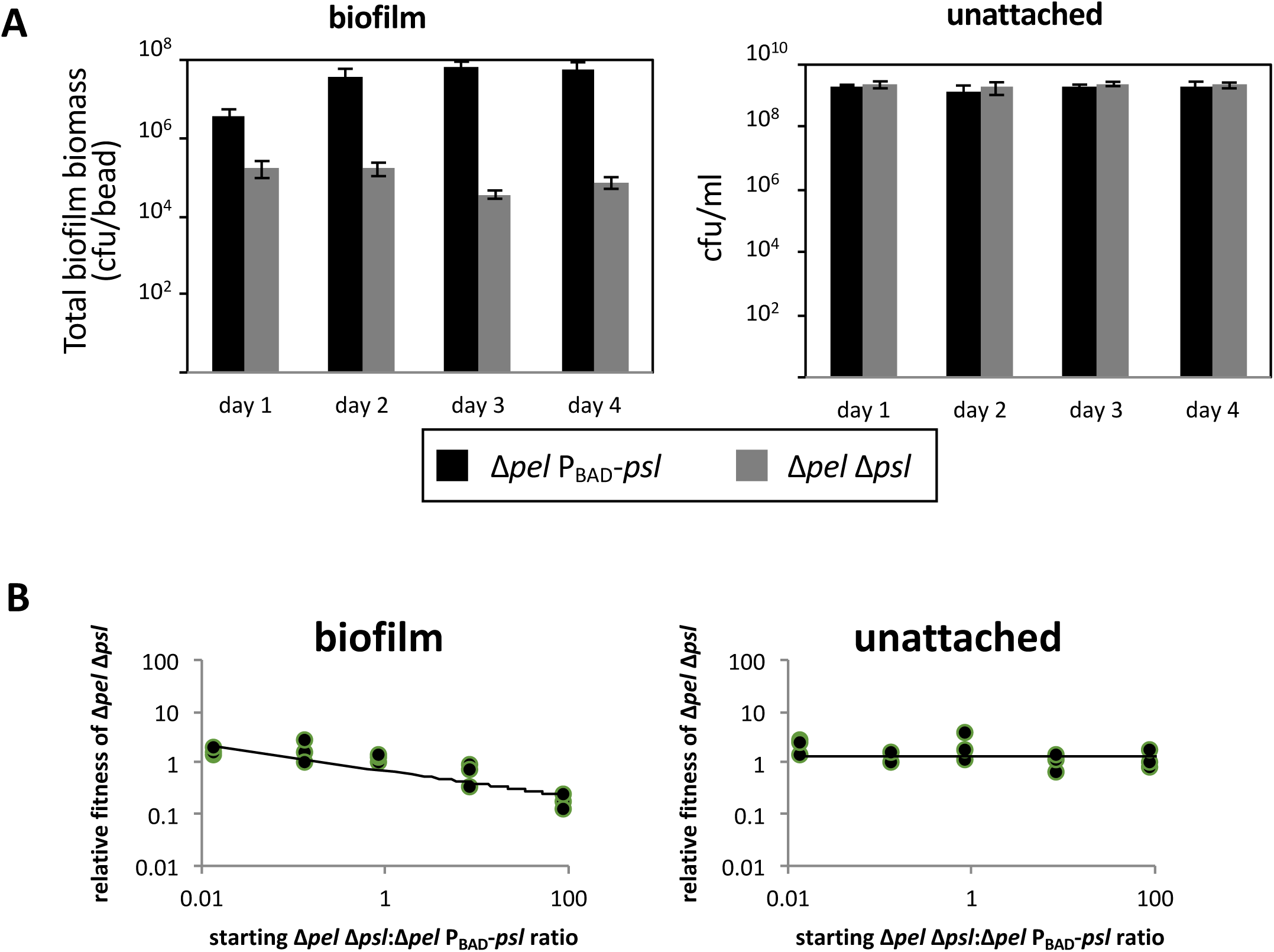
PSL-dependent social traits are independent of c-di-GMP. (*A*) Arabinose-induced PSL over-expressing strain Δ*pel* P_BAD_-*psl* consistently produced more biofilm biomass compared to the defective mutant Δ*pel* Δ*psl* (F_(1,16)_ = 17.54, *p =* 0.0007). Conversely, no significant differences between the strains were seen in the unattached fraction of the cultures (F_(1,15)_ = 11.3, *p* > 0.4). (*B*) While the experiments in the main figures were performed using Δ*wspF* strain background (constitutively elevated c-di-GMP) to remove c-di-GMP and other c-di-GMP-regulated factors from playing a role in our interpretations, repeating the experiment without mutating *wspF* yielded identical results for both biofilm populations (left panel) and unattached populations (right panel) with the results shown in Fig. 2*A*, despite Δ*pel* Δ*psl* having low intracellular c-di-GMP and Δ*pel* P_BAD_-*psl* having high intracellular c-di-GMP ^5^. The cultures were grown in 1% L-arabinose.

**Fig. S3.**
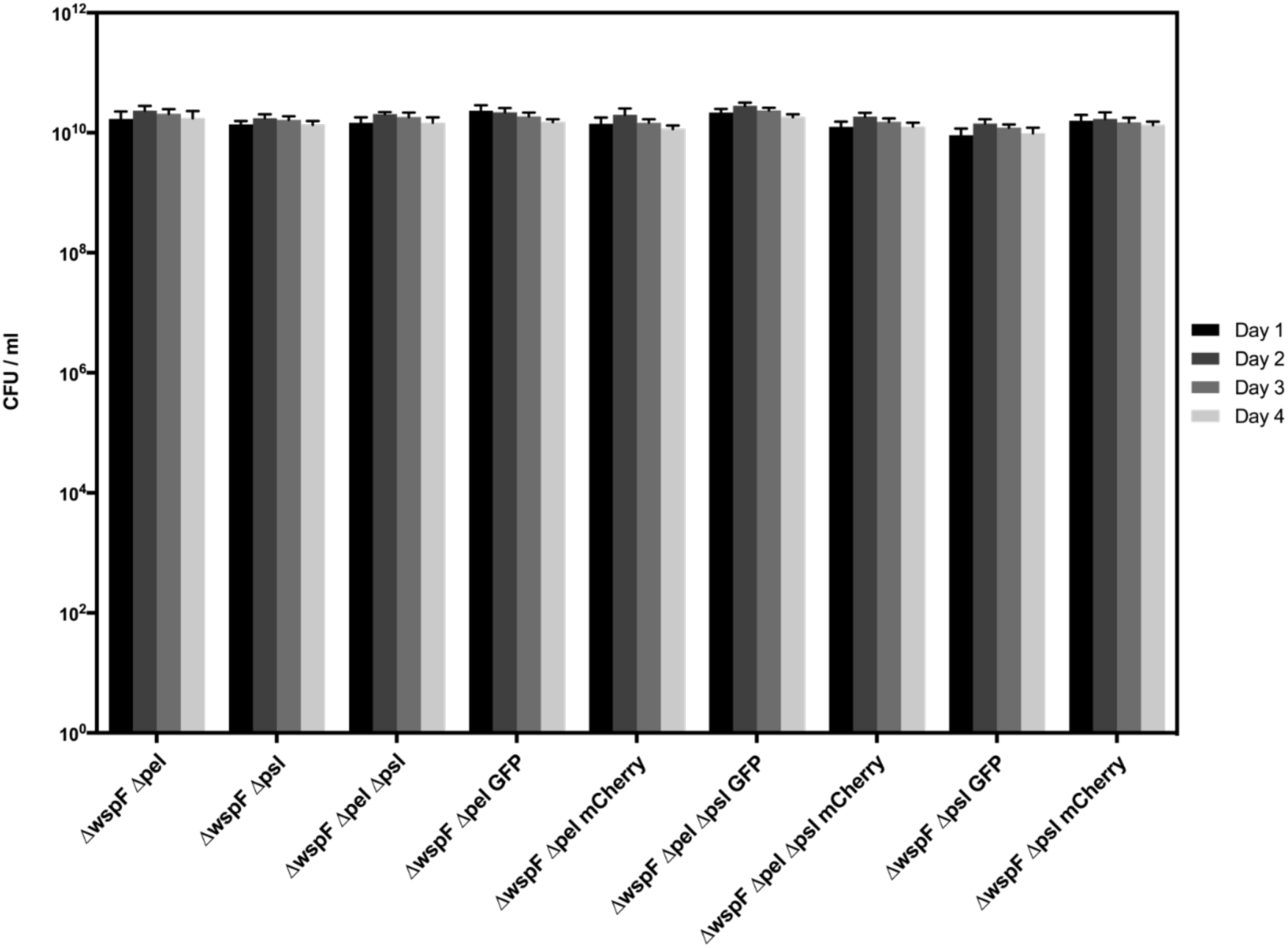
EPS mutant strains show no observable differences in growth over four-days in bead-less culture tubes.

**Fig. S4.**
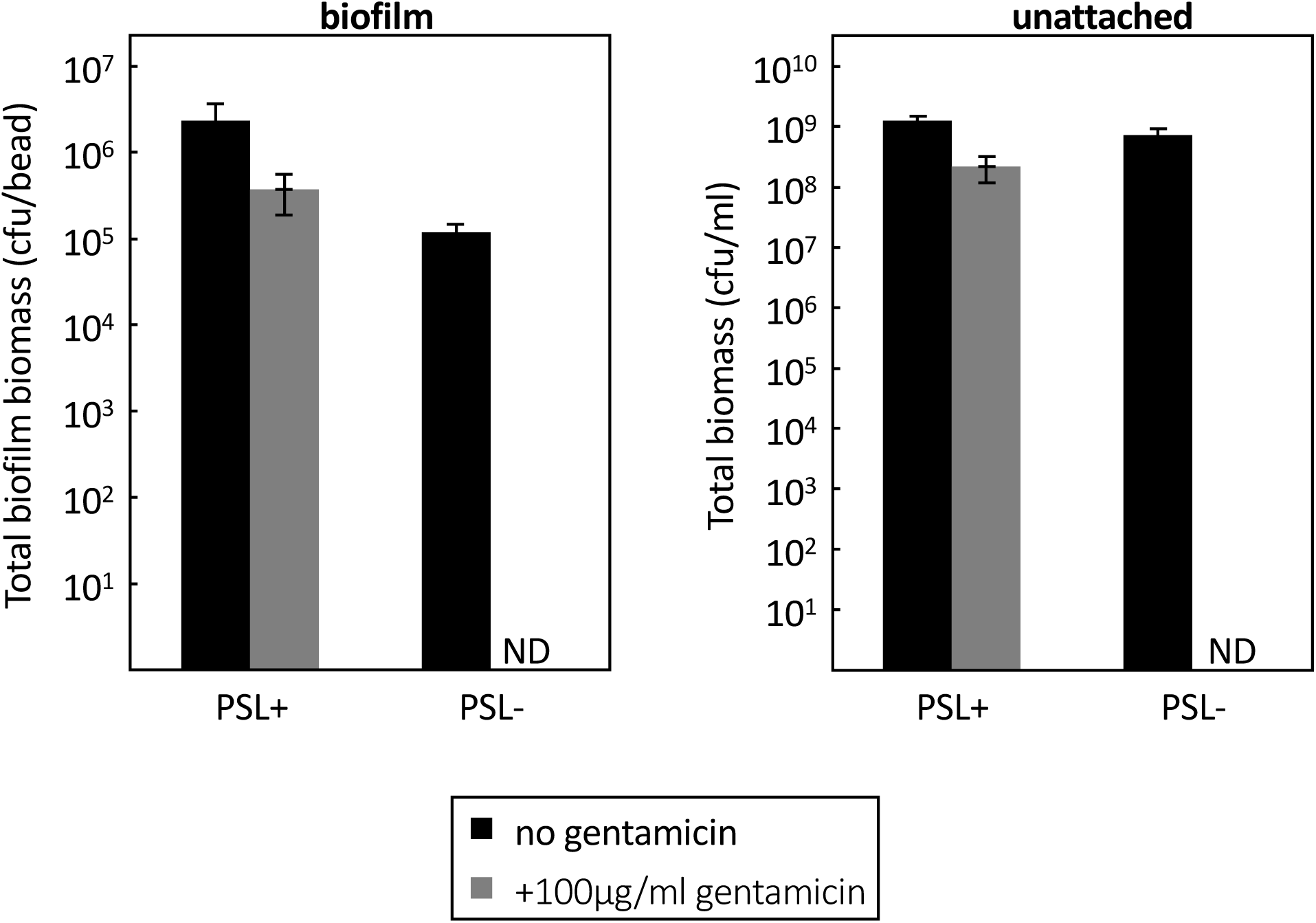
PSL expression protects cells from bactericidal concentration of gentamicin. 100 μg/ml gentamicin was added directly to 24 hours bead biofilm cultures and incubated for an additional 24 hours at 37°C with 200 rpm shaking. Resulting populations from both biofilm and unattached cells were tested for viability by serial dilution and plating. ND = not detected.

**Fig. S5.**
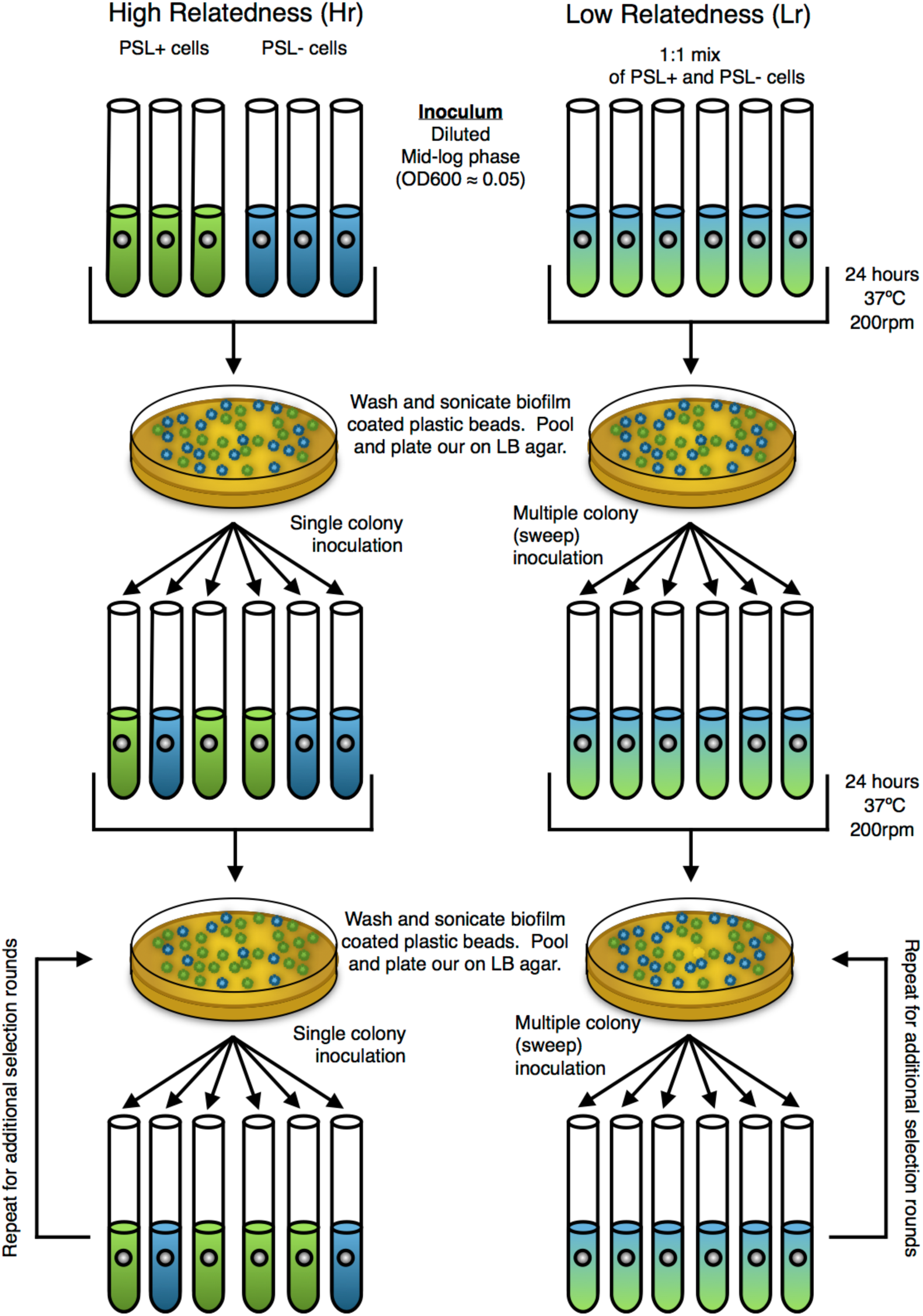
Schematics of the relatedness experiment. Tubes (populations) were initially inoculated with either single or multiple colonies (equating to populations that had relatively high relatedness [Hr] and relatively low relatedness [Lr], respectively) consisting of PSL+ cells, PSL-cells, or a 1:1 mixture of each. Populations were grown separately in the presence/absence of plastic beads and pooled together before the ratio of PSL+/− cells were determined. Single (Hr) or multiple (Lr) colonies were selected for progression into the next round of the experiment. Should one sub-population (PSL+ or PSL-) have increased fitness, the ratio and thus their likelihood to progress into subsequent rounds of selection will increase.

**Fig. S6.**
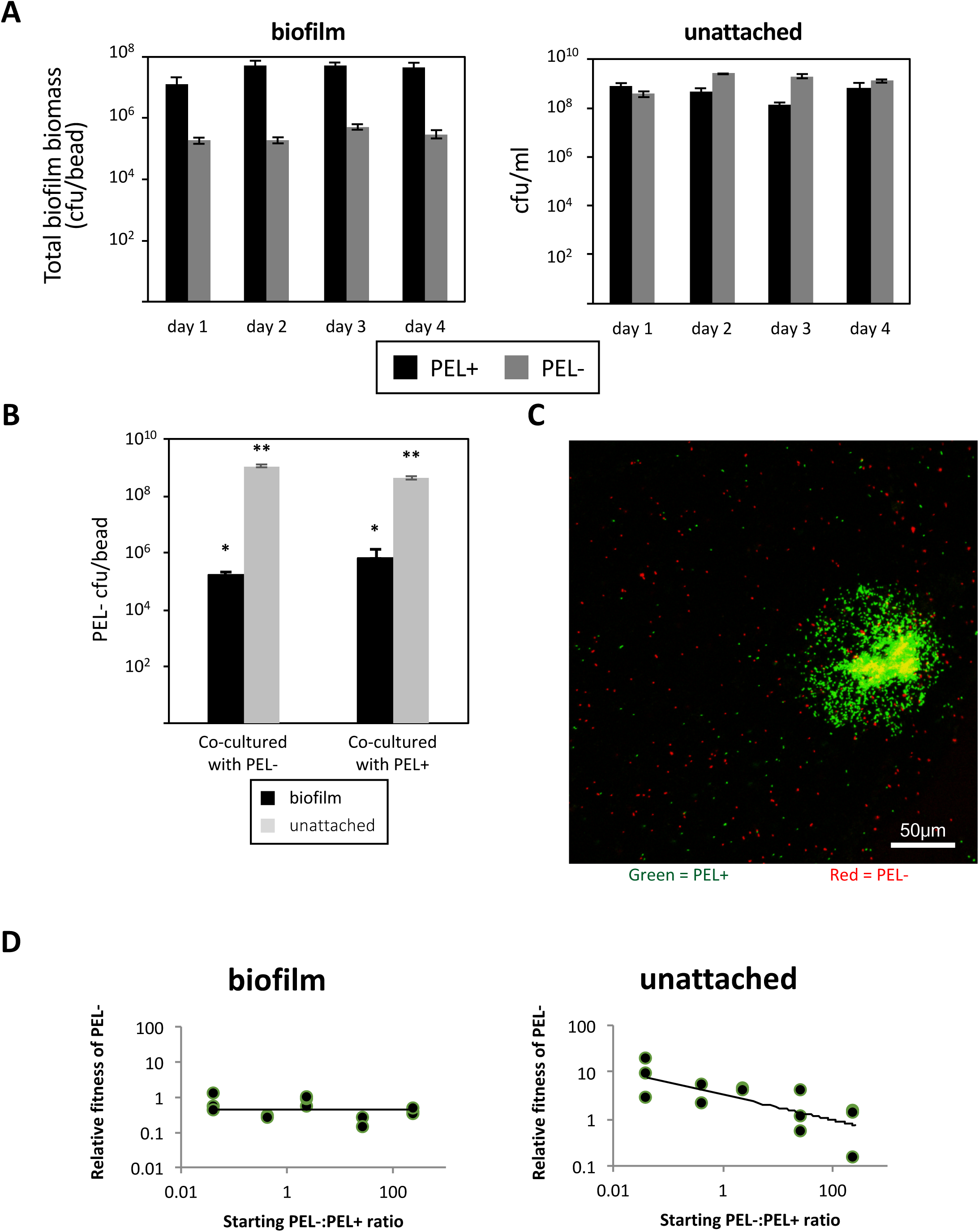
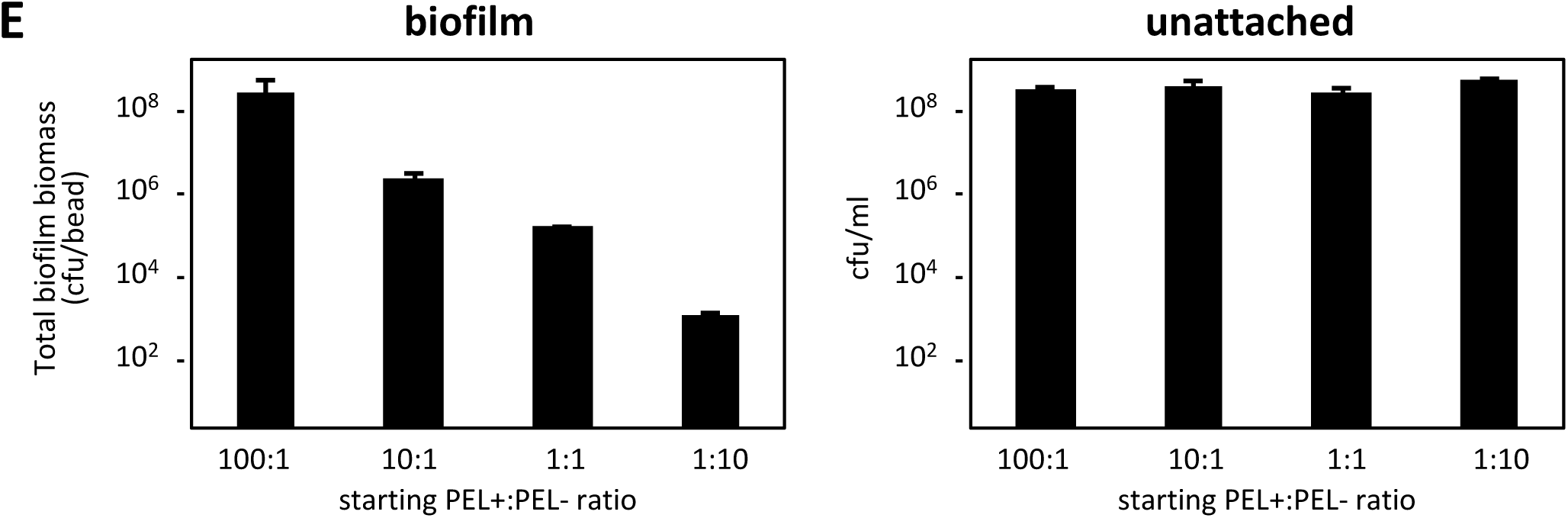
PEL polysaccharides are private goods. (*A*) Similar to PSL (Fig. 1 A), PEL-strain is significantly defective in biofilm formation compared to PEL+ (*F*_(1,24)_ = 27.3, *p* < 0.0001, *n* = 4), but no major differences are seen between their growths in the unattached fractions (*F*_(1,24)_ = 46.56, *p* < 0.0001, *n* = 4. (*B*) Unlike PSL, co-cultures of PEL-strains with PEL+ strains do not increase the amount of PEL-strains in neither biofilm nor unattached populations, signifying the unavailability of PEL polysaccharides to non-producing cells. * and ** *p >* 0.02. (*C*) Confocal micrograph image of surface attached populations of PEL-/PEL+ 1:1 co-cultures. PEL-cells (red) do not co-aggregate with PEL+ cells (green). (*D*) There is no frequency-dependent fitness changes for PEL-/PEL+ co-cultured biofilm (*t*(13) = −0.802, *p* = 0.4371), and the relative fitness is consistently slightly below 1, indicating a steady disadvantage of not expressing PEL in the biofilm. There is a frequency-dependent fitness changes in the unattached population (*t*(13) = −1.435,*p* = 0.175), perhaps caused by complex regulatory system of PEL and the involvement of quorum sensing ^8^. (*E*) Due to PEL not socially affecting co-culture communities, there is a steady decline of biofilm biomass as PEL+ becomes rarer (*F*_(1,8)_ = 1, *p* = 0.441099, *n* = 3), but no change is seen in the maximum cell numbers in the unattached population (*F*_(1,8)_ = 1.46, *p* = 0.296592, *n* = 3) regardless of the starting ratios of the strains.

